# The *E. coli* Cas1/2 endonuclease complex reduces CRISPR/Cascade guide array stability

**DOI:** 10.1101/2020.07.27.223214

**Authors:** Zhixia Ye, Juliana Lebeau, Eirik A. Moreb, Romel Menacho-Melgar, Michael D. Lynch

## Abstract

CRISPR based interference has become common in various applications from genetic circuits to dynamic metabolic control. In *E. coli* the native CRISPR Cascade system can be utilized for silencing by deletion of the *cas3* nuclease along with expression of guide RNA arrays, where multiple genes can be silenced from a single transcript. We notice the loss of protospacer sequences from guide arrays utilized for dynamic silencing. We report that unstable guide arrays are due to expression of the Cas1/2 endonuclease complex. A *cas1* deletion improves guide array stability. We propose a model wherein basal Cas1/2 endonuclease activity results in the loss of protospacers from guide arrays. Subsequently, mutant guide arrays can be amplified through selection. Replacing a constitutive promoter driving Cascade complex expression with a tightly controlled inducible promoter improves guide array stability, while minimizing leaky gene silencing.

**Highlights:**

- Cas1/2 endonuclease complex mediates CRISPR/Cascade protospacer loss in *E. coli*
- Tightly controlled Cascade operon expression increases guide array stability.

## Introduction

Gene silencing is a powerful tool and CRISPR based methods have increased the simplicity of this approach.^1^ In *E. coli*, the native multi-protein Cascade (type I-E CRISPR) system can be engineered for use in gene silencing, which involves deletion of the nuclease component and overexpression of the genes responsible for processing CRISPR arrays and target DNA binding. ^2–5^ One benefit of using the modified Cascade system is the targeting of multiple genes with the expression of a single transcript containing multiple protospacers, which is subsequently processed into individual guide RNAs. ^2^ This system enables silencing of multiple genes with rapidly constructed guide RNA arrays, and can be used in metabolic engineering strategies relying on two-stage dynamic metabolic control.^6,7^ In this approach, levels of metabolic enzymes are dynamically reduced during a phosphate depleted stationary phase cultures by a combination of Cascade based gene silencing and controlled proteolysis.^8,9^ Controlled proteolysis is implemented through the use of C-terminal DAS+4 (DAS4) degron tags, which target a given protein to the ClpXP protease only when the SspB chaperone is expressed. ^10,11^

We previously noted that silencing plasmids containing guide arrays had stability issues. ^9,12^ This instability necessitated a PCR based quality control check on strains constructed with guide array plasmids, and in several cases stable strains were not identified for certain combinations of host strains and guide RNA arrays and had to be removed from our studies. ^12^In this work, we investigate the cause of this guide array instability, with a goal toward engineering improved silencing.

## Results

Toward this aim we first sequenced several guide array plasmids where guide loss was suspected. As an example, we transformed a single guide array plasmid containing protospacers to silence the *gltAp1* (“g1”), *gltAp2* (“g2”) and *udhA* (“u”) promoters, into a host strain (DLF_Z0047) engineered with degron tags capable of proteolytic degradation of FabI (enoyl-ACP reductase), GltA (citrate synthase) and UdhA (soluble transhydrogenase). A single colony was chosen and used to inoculate a 5 mL culture (Luria broth) and, after overnight growth, the culture was plated to isolate single colonies. 24 clones were isolated and the guide array plasmid was purified and sequenced. While 17 plasmids had the expected sequence and retained all 3 protospacers (Figure 2a), the other 7 had mutations, with the loss of the 2 protospacers (“g1” and “u”) flanking the middle protospacer “g2”. Four of these modified clones retained both the 5’ and 3’ flanking repeat sequences, whereas the other three also lost either the 5’ or 3’ repeat sequence flanking the “g2” protospacer.

**Figure 1:**
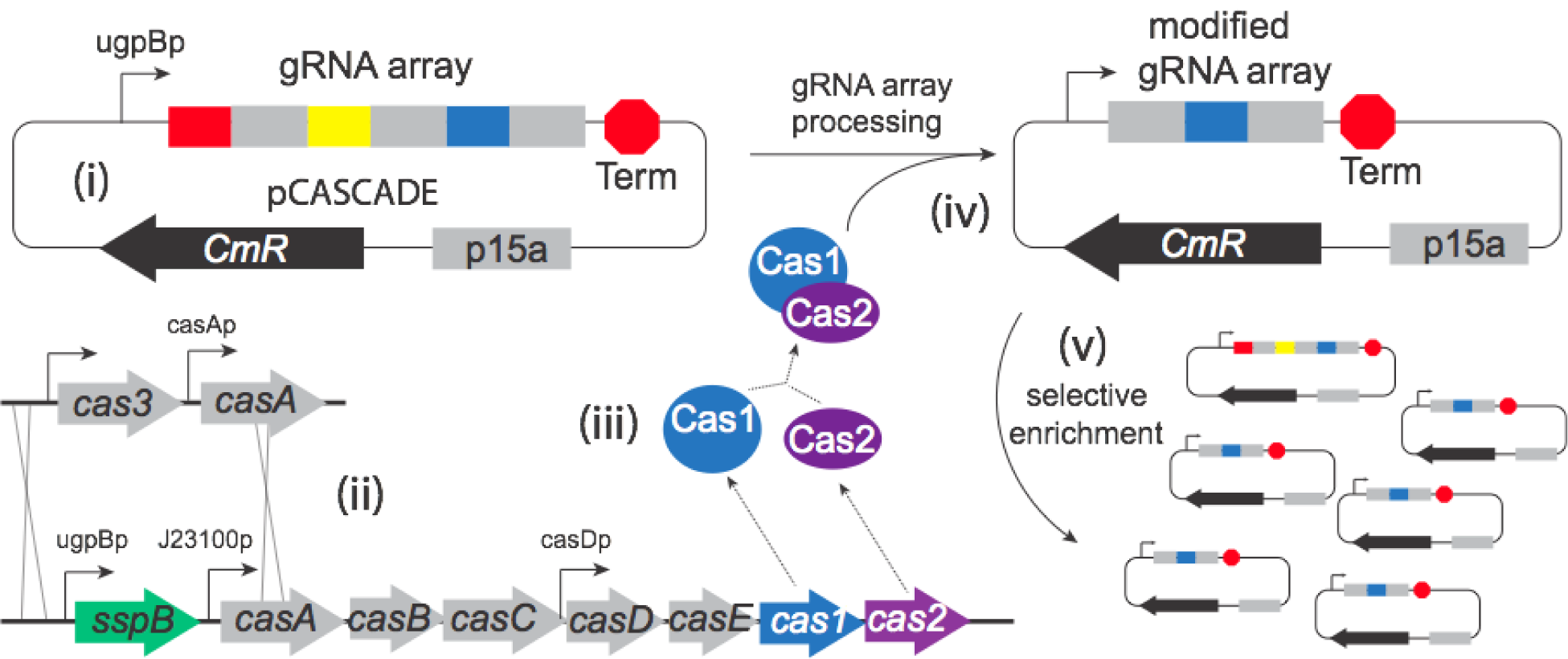
(a) Cas1/2 endonuclease mediated guide array instability. Silencing uses (i) guide arrays with multiple protospacers (colored bars, repeat sequences are in gray) as well as engineered strains where the (ii) cas3 nuclease is deleted and the cascade operon overexpressed *via* a strong constitutive promoter. (iii) Cascade operon overexpression leads to the production of the Cas1/2 endonuclease which removes (iv) protospacers from guide arrays. (v) Modified guide arrays lacking silencing capabilities are amplified through selection.

**Figure 2:**
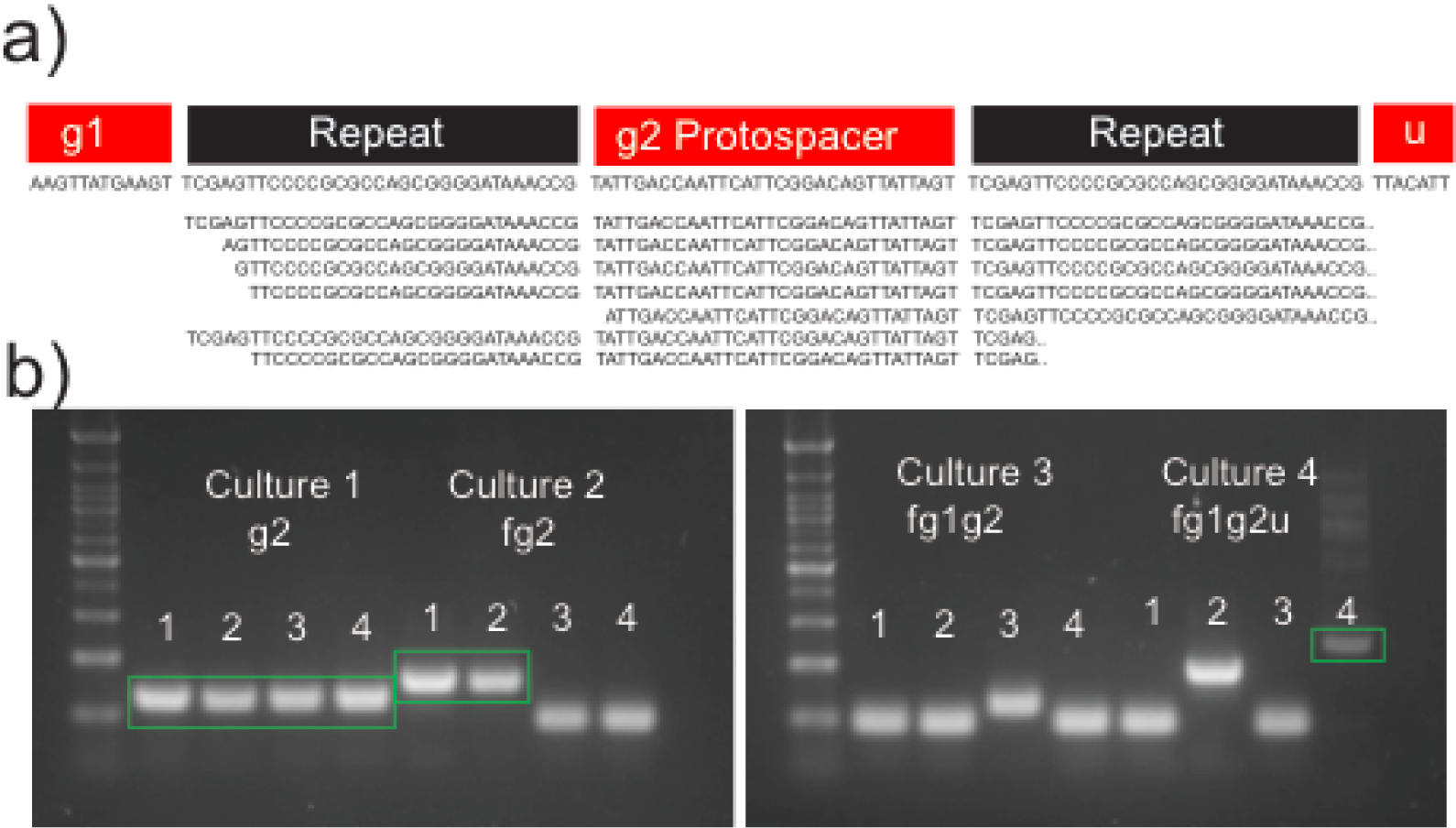
Example guide array protospacer loss. (a) Protospacer loss was noted with a guide array containing protospacers targeting the gltAp1(G1), gltAp2(G2) and udhA(U) promoters. Sequencing of modified guide arrays reveal loss of protospacer and/or repeat sequences. At the top is the correct sequence while the sequences below show results from 7 isolated clones with protospacer loss. (b) Protospacer modification can be quantified by PCR. Isolation streaking was performed with four cultures with different guide arrays and single colonies isolated for guide array PCR. Top bands highlighted in green boxes are of the expected correct size.

We next sought to evaluate the stability of a single guide (“g2”) and three additional guide arrays: “fg2”, “fg1g2” and “fg1g2u”, where “f” is a protospacer targeting the *fabI* promoter. Using the same strain, DLF_Z0047, single colonies were isolated following the same protocol described above. In this case, four clones from each of four cultures were isolated and colony PCR rather than sequencing was utilized to evaluate guide array stability. Results are given in Figure 2b. While in this case the single “g2” guide proved stable, the larger arrays of 2-4 protospacers had varying degrees of instability, producing amplicons consistent with the loss of 1-3 protospacers.

With the success of PCR as a tool to assess stability, we evaluated guide array stability for a larger grouping of guide arrays in several different host strains. These included the “f”, “g1”, “g2” and “u” protospacers as well as a protospacer targeting the *zwf* promoter, “z”. Strains that were evaluated included *E. cloni* 10G, a commercial recA1 cloning strain (Lucigen), as well as DLF_Z0025, a control host utilized for 2-stage dynamic metabolic control that lacks proteolytic degron tags on any metabolic enzymes, DLF_Z0045, with degron tags on GltA and UdhA, DLF_Z0047 (FGU, described above), as well as derivatives of DLF_Z0047 including a recA1 mutant (recAG160D), an *sbcD* gene deletion (a component of the SbcCD endonuclease recognizing hairpins and palindromic sequences present in guide arrays) and deletions in *cas1* and *cas2*.^9,13–23^ Results are given in Figure 3. Guide arrays were stable in the cloning strain. This result was not surprising as these constructs were originally constructed using E. cloni 10G and original plasmids confirmed via sequencing without any note of any protospacer loss. Protospacer loss was first noticed in DLF_Z0025 for a small group of arrays. DLF_Z0025 has been modified for constitutive expression of the Cascade operon (Figure 1). Increased instability was detected with host strain DLF_Z0045 and DLF_Z0047. Neither incorporation of a recA1 mutation nor deletion of *sbcD* reduced protospacer loss in the DLF_Z0047 background. In the case of recA, this is consistent with previous studies demonstrating that although only 20 bp of homology will enable recombination in *E. coli*, homologous sequences greater than 50bp are required for significant recombination; protospacers are 30 bp long.^24^ In contrast, both a deletion of *cas1* or *cas2* improved array stability, with a *cas1* deletion having minimal protospacer loss.

**Figure 3:**
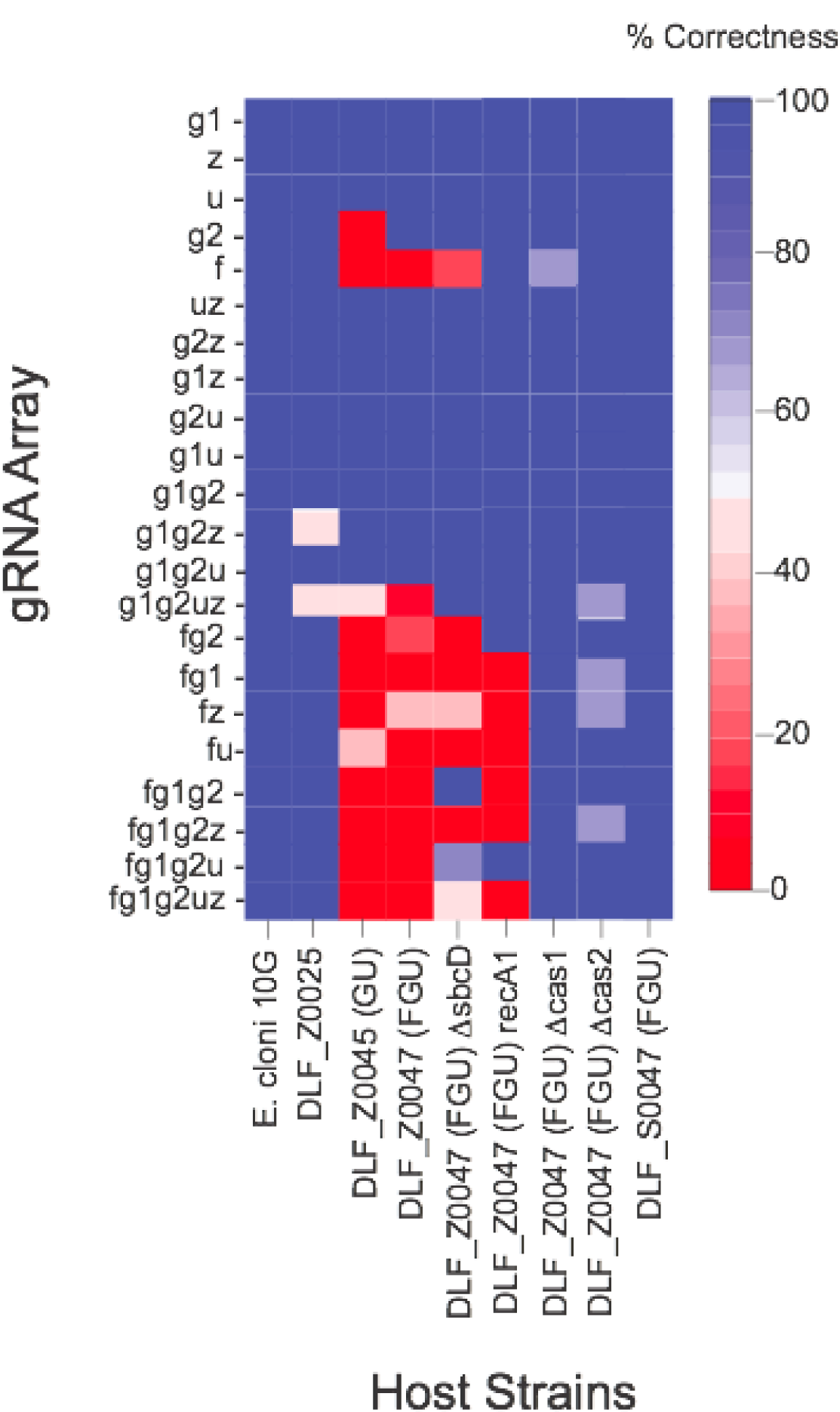
Guide array stability as a function of guide array and host strains. % correctness is the number of colony PCRs of the expected size over the total number of colonies screened. For each strain a minimum of four and maximum of twelve clones were evaluated by PCR

The *cas1* deletion results are consistent with the Cas1/2 endonuclease being responsible for protospacer loss, Cas1 being the nuclease component. This activity is consistent with their previously reported activity in protospacer acquisition.^20,22,23,25^ The fact that a very low-level of protospacer loss was still observed with a Cas1 mutation indicates the potential for a second alternative mechanism for protospacer loss, or alternatively inaccuracies in our PCR assay. However, as can be seen in Figure 3, guide arrays containing the “f” protospacer had noticeably more instability than those without. This protospacer specificity is not consistent with a generalized endonuclease activity, prompting us to further investigate why “f” containing arrays have an increased propensity for protospacer loss.

We hypothesized that as *fabI* is a strictly essential gene, ^26–28^ and despite the fact that the guide arrays are under inducible expression, even low levels of leaky expression could lead to growth inhibition, thereby giving a selective advantage to guide arrays losing the “f” protospacer in strains where the Cascade operon (including *cas1* and *cas2*) is overexpressed. This is also consistent with a general anecdotal observation in our lab that transformation of guide array plasmids with an “f” protospacer results in lower colony numbers than other arrays. In order to test this hypothesis, we constructed a plasmid (pFABI, Figure 4a) enabling the expression of FabI from an alternative constitutive EM7 promoter, which is not silenced by the “f” protospacer. We then assessed the impact of cotransformation of pFABI with guide array plasmids containing “f” protospacers on colony numbers as well as array stability. As can be seen in Figure 4 (b and c), cotransformation of pFABI increased colony numbers as well as array stability. These data are consistent with growth inhibition from leaky silencing of *fabI* giving a selective advantage to arrays where the “f” protospacer is lost.

**Figure 4:**
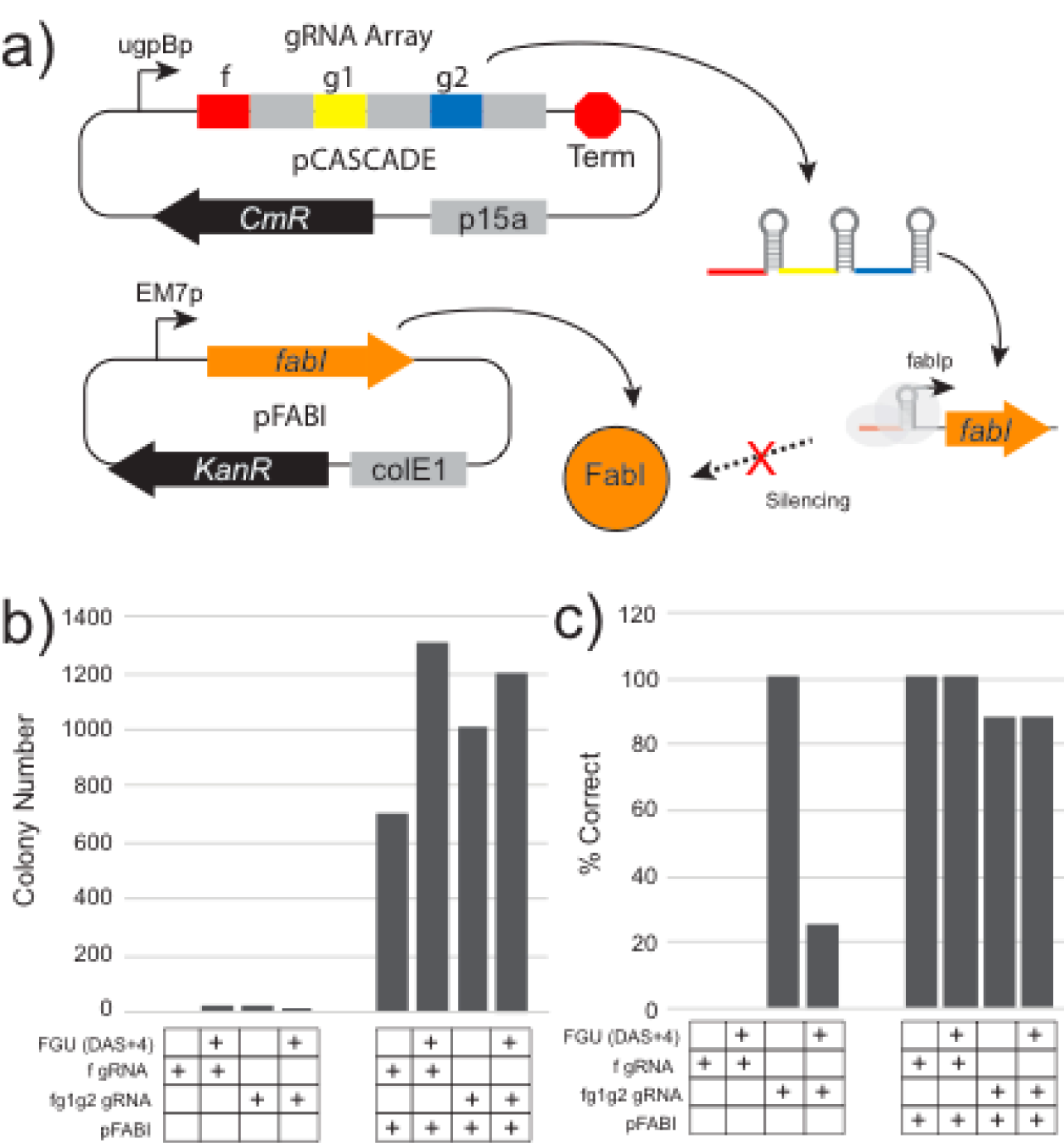
Complementation of *fabI* silencing with pFABI, a plasmid enabling overexpression of the *fabI* gene driven by a different constitutive promoter. b) Colony counts and c) guide array stability with strains transformed with guide arrays as well as pFABI.

Taken together, these support a model wherein basal Cas1/2 endonuclease activity results in the loss of protospacers from guide arrays. Silencing arrays with protospacers targeting essential genes may lead to growth inhibition, even if subtle, due to leaky expression of guides when the Cascade operon is overexpressed. Arrays missing toxic protospacers can be amplified via selection in routine cultures. In light of this understanding there are several options to improve array stability. Firstly, simply deleting *cas1* should improve stability. As Cas1 is not required for the silencing function of the Cascade operon, gene silencing should not be affected. ^3,29^ This approach would require two modifications to future silencing strains, the deletion of *cas3* and *cas1* (Figure 1a). However, in light of the toxicity observed in case of basal *fabI* silencing, we opted to evaluate a second option, wherein we deleted *cas3* and used a tightly controlled low phosphate inducible promoter to express the Cascade operon rather than a constitutive promoter (Biobrick J23100) as originally reported. ^2,30^ To implement and test this approach we constructed strain DLF_S0047, identical to DLF_Z0047, containing degron tags on FabI, GltA and UdhA, but wherein the constitutive J23100 promoter (Figure 1a) was replaced by a tightly controlled low phosphate inducible modified *yibD* gene promoter, ^31^ preceded by a strong synthetic transcriptional tZ terminator.^32,33^ Array stability was improved using DLF_S0047 as can be seen in Figure 3. With this success, we also constructed DLF_S0025 as a new stable strain for future engineering for dynamic metabolic control.

Initially, protospacer loss was only identified due to unexpectedly large errors in strain evaluations as well as inconsistent issues with strain growth. These results highlight the importance of quality control in strain construction and experimentation in synthetic biology. Future utilization of Cascade for CRISPR interference will benefit from tighter control over Cascade operon (*cas1/2*) expression, if not deletion of *cas1*, or at least evaluation of guide stability. In addition, these results support the importance of the Cas1/2 endonuclease in protospacer loss and/or modification, a central step in bacterial immunity, requiring further study.

**Table 1:**
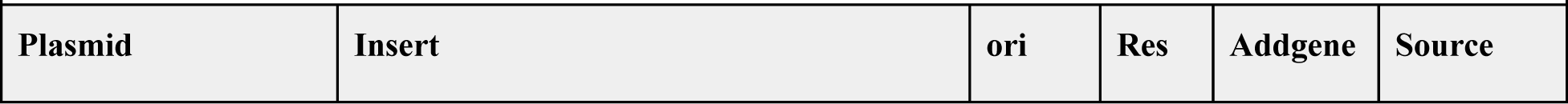
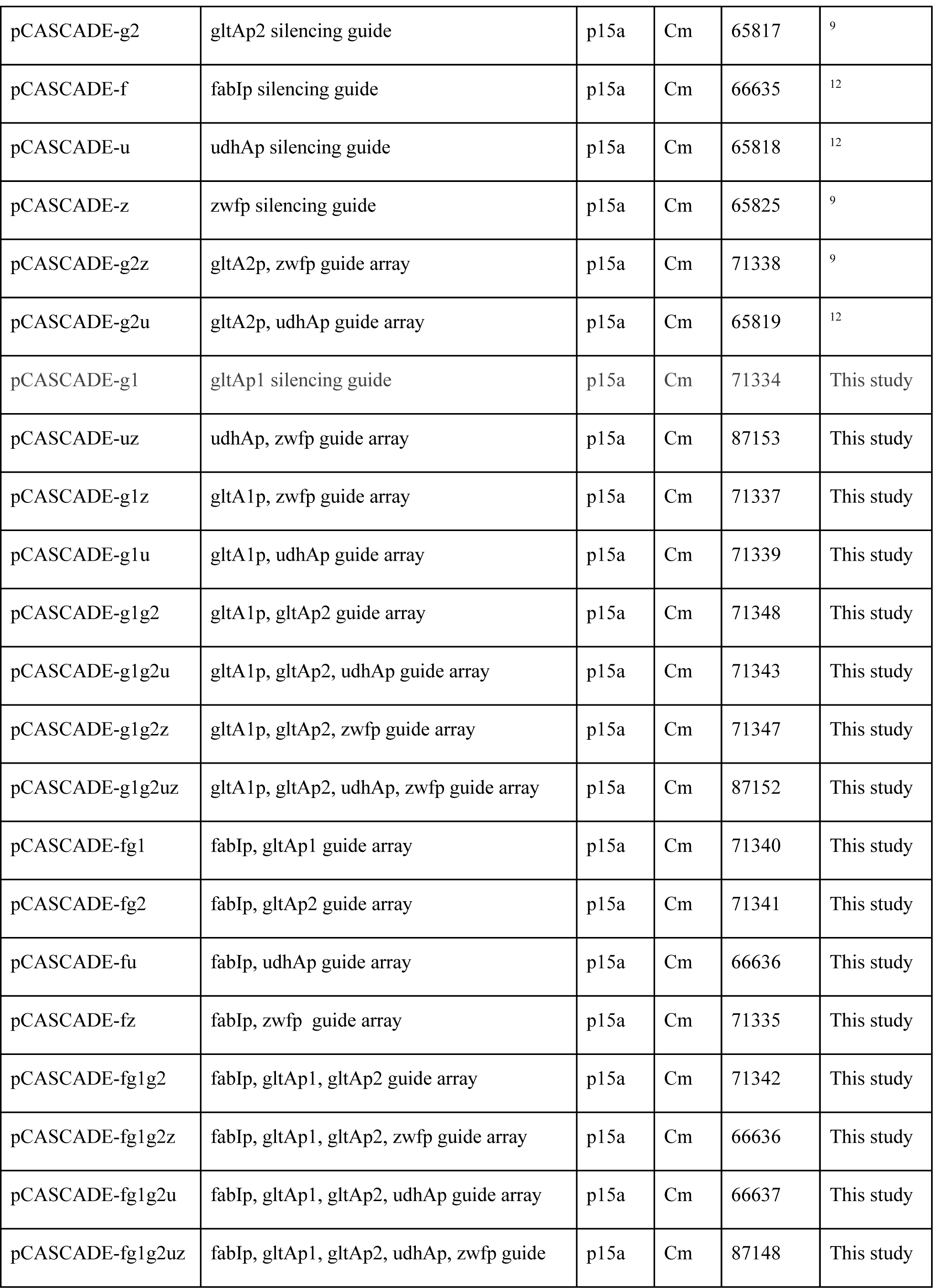
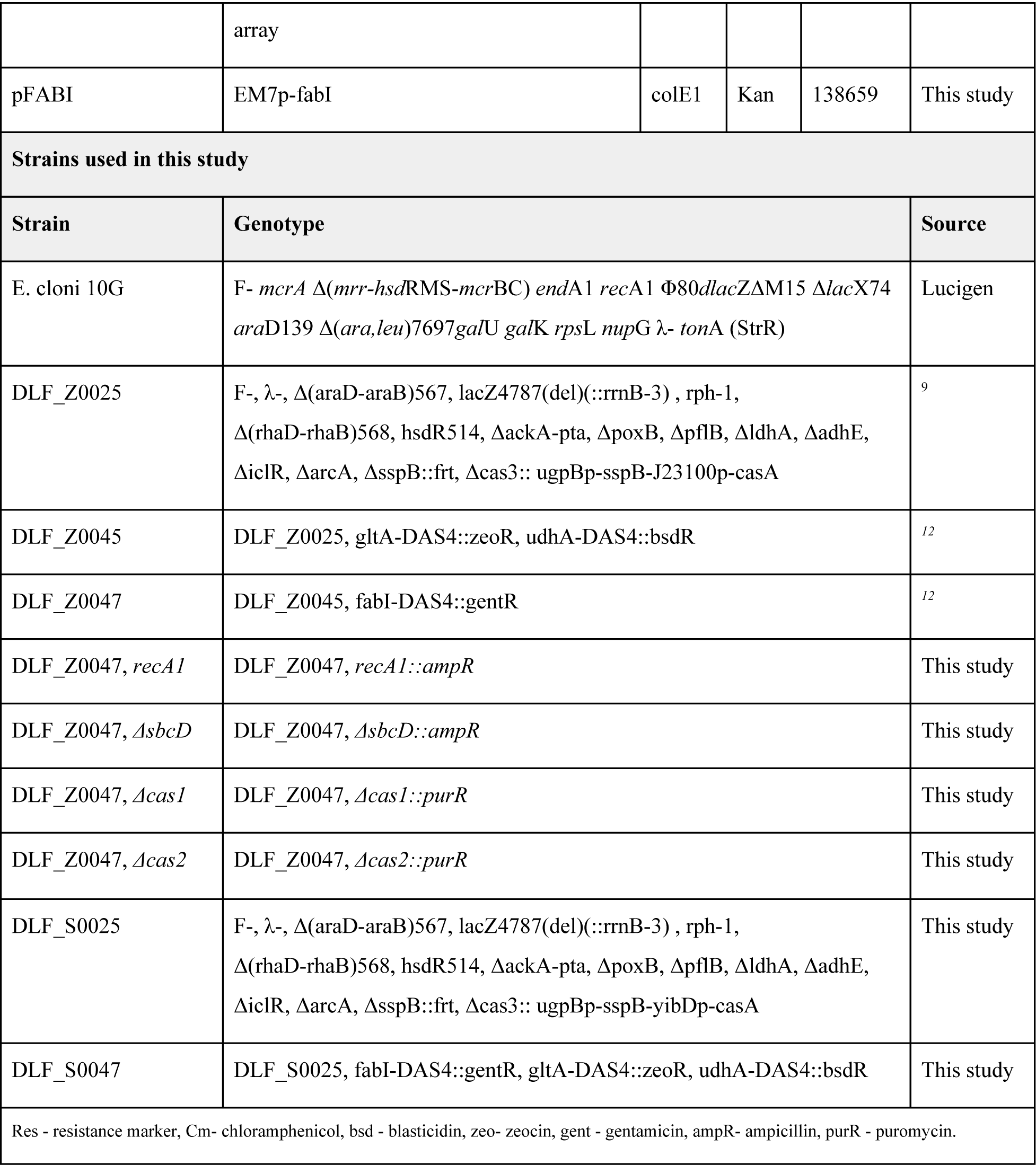
Plasmids and strain used in this study.

## Materials & Methods

### Reagents and Media

Unless otherwise stated, all materials and reagents were of the highest grade possible and purchased from Sigma (St. Louis, MO). Luria Broth, lennox formulation with lower salt was used for routine strain and plasmid propagation, construction and colony isolation. Chloramphenicol, ampicillin, and tetracycline were used at a final working concentration of 35 µg/mL, 100µg/mL, and 5 µg/mL respectively. Puromycin selection was performed using a final working concentration of 150 µg/mL, with LB supplemented with 50 mM potassium phosphate buffer (pH=8.0) to maintain pH for adequate selection.

### Strains and Plasmids

pCASCADE array plasmids were constructed as previously reported using PCR to exchange protospacers and PCR and Gibson assembly to build larger arrays from smaller arrays. ^9^ For pCASCADE plasmids constructed in this study, refer to Supplemental Materials for sequence and primer details. Plasmid, pFABI, was constructed to enable constitutive expression from a codon optimized *fabI* gene using the strong synthetic EM7 promoter. Plasmid DNA containing the promoter and gene was obtained from Twist Biosciences (San Francisco, CA). Strain *E. cloni* 10G was obtained from Lucigen. Strains DLF_Z0025, DLF_Z0045 and DLF_Z0047 were made as previously reported. All strains made in this study were constructed using standard recombineering. The recombineering plasmid pSIM5 and the tet-sacB selection/counterselection marker cassette were kind gifts from Donald Court (NCI, https://redrecombineering.ncifcrf.gov/court-lab.html). ^34,35^ Refer to Supplemental Materials for linear donor DNA sequences. DLF_Z0047 *ΔsbcD::ampR* was constructed *via* direct integration and gene replacement with linear donor DNA containing the appropriate antibiotic marker. The donor was prepared by PCR of synthetic ampicillin resistance cassette (ampR2) with primer del_sbcD_p1 and del_sbcD_p2. DLF_Z0047, *recA1::ampR* was similarly constructed, however the integration incorporated a G160D mutation into the *recA* gene rather than a deletion. Strains DLF_Z0047 *Δcas1::purR* and DLF_Z0047 *Δcas2::purR* were constructed *via* direct integration and gene replacement with linear donor DNA. Strains DLF_S0047 and DLF_S0025 were constructed from DLF_Z0047 and DLF_Z0025 respectively, using recombineering and tet-sacB based selection counterselection to replace the *sspB* gene and promoter in front of the Cascade operon. All genetic modifications were confirmed by PCR and sequencing. Sequencing was performed by either Genewiz (Morrisville, NC) or Eurofins (Louisville, KY). Plasmid transformations were accomplished using standard methods.

### Guide Stability Testing

Plasmid DNA minipreps and sequencing were performed with standard methods. The following two primers were used to amplify guide arrays from pCASCADE plasmids gRNA-for: 5’-GGGAGACCACAACGG-3’, gRNA-rev: 5’-CGCAGTCGAACGACCG-3’. Colony PCR was performed as follows: 2X EconoTaq Master mix (Lucigen) was used in 20 µL PCR reactions consisting of 10µL of 2X EconoTaq Master mix (Lucigen), 1uL of each primer (10uM concentration), 8uL dH2O and a small part of a colony. PCR parameters were: an initial 98°C, 2 minute initial denaturation followed by 35 cycles of 94°C, 30 seconds, 50°C 30 seconds, and 72°C, 30 seconds and a final 72°C, 5 min final extension. PCR products were then analyzed via 2% agarose gel electrophoresis.

## Acknowledgements

We would like to acknowledge support from: ONR YIP #12043956, and DOE EERE grant #EE0007563, as well as support from DMC Biotechnologies, Inc. We would also like to acknowledge M. Maurer for his help in constructing strains DLF_Z0047, *recA1::ampR* and DLF_Z0047, *ΔsbcD::ampR*.

## Author contributions

Z. Ye and R. Menacho-Melgar constructed strains and plasmids. Z. Ye and J. Lebeau performed transformations and stability assessments. E.A. Moreb contributed to hypotheses and experimental designs. M.D. Lynch designed and analyzed experimental results. All authors wrote, revised and edited the manuscript.

