## Supplementary Materials for "The *E. coli* Cas1/2 endonuclease complex reduces CRISPR/Cascade guide array stability"

**Supplemental Materials**

**Table S1:** List of sgRNA guide sequences and primers used to construct them. Spacers are italicized.

| **sgRNA/Primer Name** | **Sequence** | **Template** |
|---|---|---|
| gltA1 | *TCGAGTTCCCCGCGCCAGCGGGGATAAACCG*AAAAGCATATAATGCGTAAAAGTTATGAAGT*TCGAGTTCCCCGCGCCAGCGGGGATAAACCG* | |
| gltA1-FOR | GCGTAAAAGTTATGAAGT*TCGAGTTCCCCGCGCCAGCGGGGATAAACCG*AAAAAAAAACCCC | pCASCADE control |
| gltA1-REV | ATTATATGCTTTTCGGTTTATCCCCGCTGGCGCGGGGAACTCGAGGTGGTACCAGATCT | |
| G1Z | *TCGAGTTCCCCGCGCCAGCGGGGATAAACCG*AAAAGCATATAATGCGTAAAAGTTATGAAGT*TCGAGTTCCCCGCGCCAGCGGGGATAAACCG*CTCGTAAAAGCAGTACAGTGCACCGTAAGA*TCGAGTTCCCCGCGCCAGCGGGGATAAACCG* | |
| zwf-FOR | *GCGCCAGCGGGGATAAACCGCTCGTAAAAG* | pCASCADE-zwf |
| pCASCADE-REV | *CTTGCCCGCCTGATGAATGCTCATCCGG* | |
| pCASCADE-FOR | *CCGGATGAGCATTCATCAGGCGGGCAAG* | pCASCADE-gltA1 |
| gltA1-REV | *CGGTTTATCCCCGCTGGCGCGGGGAACTCGAACTTCATAACTTTTAC* | |
| G1U | *TCGAGTTCCCCGCGCCAGCGGGGATAAACCG*AAAAGCATATAATGCGTAAAAGTTATGAAGT*TCGAGTTCCCCGCGCCAGCGGGGATAAACCG*TTACCATTCTGTTGCTTTTATGTATAAGAA*TCGAGTTCCCCGCGCCAGCGGGGATAAACCG* | |

| | | |
|---|---|---|
| udhA-FOR | *GCGCCAGCGGGGATAAACCGTTACCATTCTGTTG* | pCASCADE-udhA |
| pCASCADE-REV | *CTTGCCCGCCTGATGAATGCTCATCCGG* | |
| pCASCADE-FOR | *CCGGATGAGCATTCATCAGGCGGGCAAG* | pCASCADE-gltA1 |
| gltA1-REV | *CGGTTTATCCCCGCTGGCGCGGGGAACTCGAACTTCATAACTTTTAC* | |
| G1G2 | *TCGAGTTCCCCGCGCCAGCGGGGATAAACCG*AAAAGCATATAATGCGTAAAAGTTATGAAGT*TCGAGTTCCCCGCGCCAGCGGGGATAAACCG*TATTGACCAATTCATTCGGGACAGTTATTAGT*TCGAGTTCCCCGCGCCAGCGGGGATAAACCG* | |
| gltA2-FOR | *GCGCCAGCGGGGATAAACCGTATTGACCAATTCATTC* | pCASCADE-gltA2 |
| pCASCADE-REV | *CTTGCCCGCCTGATGAATGCTCATCCGG* | |
| pCASCADE-FOR | *CCGGATGAGCATTCATCAGGCGGGCAAG* | pCASCADE-gltA1 |
| gltA1-REV | *CGGTTTATCCCCGCTGGCGCGGGGAACTCGAACTTCATAACTTTTAC* | |
| G1G2U | *TCGAGTTCCCCGCGCCAGCGGGGATAAACCG*AAAAGCATATAATGCGTAAAAGTTATGAAGT*TCGAGTTCCCCGCGCCAGCGGGGATAAACCG*TATTGACCAATTCATTCGGGACAGTTATTAGT*TCGAGTTCCCCGCGCCAGCGGGGATAAACCG*TTACCATTCTGTTGCTTTTATGTATAAGAA*TCGAGTTCCCCGCGCCAGCGGGGATAAACCG* | |
| udhA-FOR | *GCGCCAGCGGGGATAAACCGTTACCATTCTGTTG* | pCASCADE-udhA |
| pCASCADE-REV | *CTTGCCCGCCTGATGAATGCTCATCCGG* | |

| | | |
|---|---|---|
| pCASCADE-FOR | *CCGGATGAGCATTCATCAGGCGGGCAAG* | pCASCADE-G1G2 |
| gltA2-REV | *CGGTTTATCCCCGCTGGCGCGGGGAACTCGAACTAATAACTGTC* | |
| G1G2Z | *TCGAGTTCCCCGCGCCAGCGGGGATAAACCG*AAAAGCATATAATGCGTAAAAGTTATGAAGT*TCGAGTTCCCCGCGCCAGCGGGGATAAACCG*TATTGACCAATTCATTCGGGACAGTTATTAGT*TCGAGTTCCCCGCGCCAGCGGGGATAAACCG*CTCGTAAAAGCAGTACAGTGCACCGTAAGA*TCGAGTTCCCCGCGCCAGCGGGGATAAACCG* | |
| zwf-FOR | *GCGCCAGCGGGGATAAACCGCTCGTAAAAG* | pCASCADE-zwf |
| pCASCADE-REV | *CTTGCCCGCCTGATGAATGCTCATCCGG* | |
| pCASCADE-FOR | *CCGGATGAGCATTCATCAGGCGGGCAAG* | pCASCADE-G1G2 |
| gltA2-REV | *CGGTTTATCCCCGCTGGCGCGGGGAACTCGAACTAATAACTGTC* | |
| G1G2UZ | *TCGAGTTCCCCGCGCCAGCGGGGATAAACCG*AAAAGCATATAATGCGTAAAAGTTATGAAGT*TCGAGTTCCCCGCGCCAGCGGGGATAAACCG*TATTGACCAATTCATTCGGGACAGTTATTAGT*TCGAGTTCCCCGCGCCAGCGGGGATAAACCG*TTACCATTCTGTTGCTTTTATGTATAAGAA*TCGAGTTCCCCGCGCCAGCGGGGATAAACCG*CTCGTAAAAGCAGTACAGTGCACCGTAAGA*TCGAGTTCCCCGCGCCAGCGGGGATAAACCG* | |
| zwf-FOR | *GCGCCAGCGGGGATAAACCGCTCGTAAAAG* | pCASCADE-zwf |
| pCASCADE-REV | *CTTGCCCGCCTGATGAATGCTCATCCGG* | |
| pCASCADE-FOR | *CCGGATGAGCATTCATCAGGCGGGCAAG* | pCASCADE-G1G2U |

| | | |
|---|---|---|
| udhA-REV | *CGGTTTATCCCCGCTGGCGCGGGGAACTCGATTCTTATACATAAAAGC* | |
| FG1 | *TCGAGTTCCCCGCGCCAGCGGGGATAAACCG*TTGATTATAATAACCGTTTATCTGTTCGTA*TCGAGTTCCCCGCGCCAGCGGGGATAAACCG*AAAAGCATATAATGCGTAAAAGTTATGAAGT*TCGAGTTCCCCGCGCCAGCGGGGATAAACCG* | |
| gltA1-FOR | GCGCCAGCGGGGATAAACCG*AAAAGCATATAAT*GCG | pCASCADE-gltA1 |
| pCASCADE-REV | CTTGCCCGCCTGATGAATGCTCATCCGG | |
| pCASCADE-FOR | CCGGATGAGCATTCATCAGGCGGGCAAG | pCASCADE-fabI |
| fabI-REV | CGGTTTATCCCCGCTGGCGCGGGGAACTCGATACGAACAGATAAACGGTTATTATAATC | |
| FG2 | *TCGAGTTCCCCGCGCCAGCGGGGATAAACCG*TTGATTATAATAACCGTTTATCTGTTCGTA*TCGAGTTCCCCGCGCCAGCGGGGATAAACCG*TATTGACCAATTCATTCGGGACAGTTATTAGT*TCGAGTTCCCCGCGCCAGCGGGGATAAACCG* | |
| gltA2-FOR | GCGCCAGCGGGGATAAACCG*TATTGACCAATTCATTC* | pCASCADE-gltA2 |
| pCASCADE-REV | CTTGCCCGCCTGATGAATGCTCATCCGG | |
| pCASCADE-FOR | CCGGATGAGCATTCATCAGGCGGGCAAG | pCASCADE-fabI |
| fabI-REV | CGGTTTATCCCCGCTGGCGCGGGGAACTCGATACGAACAGATAAACGGTTATTATAATC | |
| FU | *TCGAGTTCCCCGCGCCAGCGGGGATAAACCG*TTGATTATAATAACCGTTTATCTGTTCGTA*TCGAGTTCCCCGCGCCAGCGGGGATAAACCG*TTACCATTCTGTTGCTTTTATGTATAAGAA*TCGAGTTCCCCGCGCCAGCGGGGATAAACCG* | |
| udhA-FOR | GCGCCAGCGGGGATAAACCGTTACCATTCTGTTG | pCASCADE-udhA |
| pCASCADE-REV | CTTGCCCGCCTGATGAATGCTCATCCGG | |
| pCASCADE-FOR | CCGGATGAGCATTCATCAGGCGGGCAAG | pCASCADE-fabI |

| | | |
|---|---|---|
| fabI-REV | CGGTTTATCCCCGCTGGCGCGGGGAACTCGATACGAACAGATAAACGGTTATTATAATC | |
| FZ | *TCGAGTTCCCCGCGCCAGCGGGGATAAACCG*TTGATTATAATAACCGTTTATCTGTTCGTA*TCGAGTTCCCCGCGCCAGCGGGGATAAACCG*CTCGTAAAAGCAGTACAGTGCACCGTAAGA*TCGAGTTCCCCGCGCCAGCGGGGATAAACCG* | |
| zwf-FOR | GCGCCAGCGGGGATAAACCGCTCGTAAAAG | pCASCADE-zwf |
| pCASCADE-REV | CTTGCCCGCCTGATGAATGCTCATCCGG | |
| pCASCADE-FOR | CCGGATGAGCATTCATCAGGCGGGCAAG | pCASCADE-fabI |
| fabI-REV | CGGTTTATCCCCGCTGGCGCGGGGAACTCGATACGAACAGATAAACGGTTATTATAATC | |
| FG1G2 | *TCGAGTTCCCCGCGCCAGCGGGGATAAACCG*TTGATTATAATAACCGTTTATCTGTTCGTA*TCGAGTTCCCCGCGCCAGCGGGGATAAACCG*AAAAGCATATAATGCGTAAAAGTTATGAAGT*TCGAGTTCCCCGCGCCAGCGGGGATAAACCG*TATTGACCAATTCATTCGGGACAGTTATTAGT*TCGAGTTCCCCGCGCCAGCGGGGATAAACCG* | |
| gltA2-FOR | GCGCCAGCGGGGATAAACCG*TATTGACCAATTCATTC* | pCASCADE-gltA2 |
| pCASCADE-REV | CTTGCCCGCCTGATGAATGCTCATCCGG | |
| pCASCADE-FOR | CCGGATGAGCATTCATCAGGCGGGCAAG | pCASCADE-FG1 |
| gltA1-REV | CGGTTTATCCCCGCTGGCGCGGGGAACTCGAACTTCATAACTTTTAC | |
| FG1G2U | *TCGAGTTCCCCGCGCCAGCGGGGATAAACCG*TTGATTATAATAACCGTTTATCTGTTCGTA*TCGAGTTCCCCGCGCCAGCGGGGATAAACCG*AAAAGCATATAATGCGTAAAAGTTATGAAGT*TCGAGTTCCCCGCGCCAGCGGGGATAAACCG*TATTGACCAATTCATTCGGGACAGTTATTAGT*TCGAGTTCCCCGCGCCAGCGGGGATAAACCG*TTACCATTCTGTTGCTTTTATGTATAAGAA*TCGAGTTCCCCGCGCCAGCGGGGATAAACCG* | |
| gltA2-FOR | GCGCCAGCGGGGATAAACCG*TATTGACCAATTCATTC* | pCASCADE-udhA |
| pCASCADE-REV | CTTGCCCGCCTGATGAATGCTCATCCGG | |

| | | |
|---|---|---|
| pCASCADE-FOR | CCGGATGAGCATTCATCAGGCGGGCAAG | pCASCADE-FG1G2 |
| gltA1-REV | CGGTTTATCCCCGCTGGCGCGGGGAACTCGAACTTCATAACTTTTAC | |
| FG1G2Z | *TCGAGTTCCCCGCGCCAGCGGGGATAAACCG*TTGATTATAATAACCGTTTATCTGTTCGTA*TCGAGTTCCCCGCGCCAGCGGGGATAAACCG*AAAAGCATATAATGCGTAAAAGTTATGAAGT*TCGAGTTCCCCGCGCCAGCGGGGATAAACCG*TATTGACCAATTCATTCGGGACAGTTATTAGT*TCGAGTTCCCCGCGCCAGCGGGGATAAACCG*CTCGTAAAAGCAGTACAGTGCACCGTAAGA*TCGAGTTCCCCGCGCCAGCGGGGATAAACCG* | |
| gltA2-FOR | GCGCCAGCGGGGATAAACCG*TATTGACCAATTCATTC* | pCASCADE-zwf |
| pCASCADE-REV | CTTGCCCGCCTGATGAATGCTCATCCGG | |
| pCASCADE-FOR | CCGGATGAGCATTCATCAGGCGGGCAAG | pCASCADE-FG1G2 |
| gltA1-REV | CGGTTTATCCCCGCTGGCGCGGGGAACTCGAACTTCATAACTTTTAC | |
| FG1G2UZ | *TCGAGTTCCCCGCGCCAGCGGGGATAAACCG*TTGATTATAATAACCGTTTATCTGTTCGTA*TCGAGTTCCCCGCGCCAGCGGGGATAAACCG*AAAAGCATATAATGCGTAAAAGTTATGAAGT*TCGAGTTCCCCGCGCCAGCGGGGATAAACCG*TATTGACCAATTCATTCGGGACAGTTATTAGT*TCGAGTTCCCCGCGCCAGCGGGGATAAACCG*TTACCATTCTGTTGCTTTTATGTATAAGAA*TCGAGTTCCCCGCGCCAGCGGGGATAAACCG*CTCGTAAAAGCAGTACAGTGCACCGTAAGA*TCGAGTTCCCCGCGCCAGCGGGGATAAACCG* | |
| zwf-FOR | GCGCCAGCGGGGATAAACCGCTCGTAAAAG | pCASCADE-zwf |
| pCASCADE-REV | CTTGCCCGCCTGATGAATGCTCATCCGG | |
| pCASCADE-FOR | CCGGATGAGCATTCATCAGGCGGGCAAG | pCASCADE-FG1G2U |
| udhA-REV | CGGTTTATCCCCGCTGGCGCGGGGAACTCGATTCTTATACATAAAAGC | |
| UZ | *TCGAGTTCCCCGCGCCAGCGGGGATAAACCG*TTACCATTCTGTTGCTTTTATGTATAAGAA*TCGAGTTCCCCGCGCCAGCGGG* | |

| | | |
|---|---|---|
| | *GATAAACCG*CTCGTAAAAGCAGTACAGTGCACCGTAAGA*T CGAGTTCCCCGCGCCAGCGGGGATAAACCG* | |
| zwf-FOR | GCGCCAGCGGGGATAAACCGCTCGTAAAAG | pCASCADE-zwf |
| pCASCADE-REV | CTTGCCCGCCTGATGAATGCTCATCCGG | |
| pCASCADE-FOR | CCGGATGAGCATTCATCAGGCGGGCAAG | pCASCADE-udhA |
| udhA-REV | CGGTTTATCCCCGCTGGCGCGGGGAACTCGATTCTTATAC ATAAAAGC | |

**Table S2:** List of synthetic DNA used in this study

| **tetA-sacB Cassette** |
|---|
| TCCTAATTTTTGTTGACACTCTATCATTGATAGAGTTATTTTACCACTCCCTATCAGTGATA GAGAAAAGTGAAATGAATAGTTCGACAAAGATCGCATTGGTAATTACGTTACTCGATGCC ATGGGGATTGGCCTTATCATGCCAGTCTTGCCAACGTTATTACGTGAATTTATTGCTTCGG AAGATATCGCTAACCACTTTGGCGTATTGCTTGCACTTTATGCGTTAATGCAGGTTATCTTT GCTCCTTGGCTTGGAAAAATGTCTGACCGATTTGGTCGGCGCCCAGTGCTGTTGTTGTCAT TAATAGGCGCATCGCTGGATTACTTATTGCTGGCTTTTTCAAGTGCGCTTTGGATGCTGTAT TTAGGCCGTTTGCTTTCAGGGATCACAGGAGCTACTGGGGCTGTCGCGGCATCGGTCATTG CCGATACCACCTCAGCTTCTCAACGCGTGAAGTGGTTCGGTTGGTTAGGGGCAAGTTTTGG GCTTGGTTTAATAGCGGGGCCTATTATTGGTGGTTTTGCAGGAGAGATTTCACCGCATAGT CCCTTTTTTATCGCTGCGTTGCTAAATATTGTCACTTTCCTTGTGGTTATGTTTTGGTTCCGT GAAACCAAAAATACACGTGATAATACAGATACCGAAGTAGGGGTTGAGACGCAATCGAA TTCGGTATACATCACTTTATTTAAAACGATGCCCATTTTGTTGATTATTTATTTTTCAGCGC AATTGATAGGCCAAATTCCCGCAACGGTGTGGGTGCTATTTACCGAAAATCGTTTTGGATG GAATAGCATGATGGTTGGCTTTTCATTAGCGGGTCTTGGTCTTTTACACTCAGTATTCCAAG CCTTTGTGGCAGGAAGAATAGCCACTAAATGGGGCGAAAAAACGGCAGTACTGCTCGGAT TTATTGCAGATAGTAGTGCATTTGCCTTTTTAGCGTTTATATCTGAAGGTTGGTTAGTTTTC CCTGTTTTAATTTTATTGGCTGGTGGTGGGATCGCTTTACCTGCATTACAGGGAGTGATGTC TATCCAAACAAAGAGTCATCAGCAAGGTGCTTTACAGGGATTATTGGTGAGCCTTACCAAT GCAACCGGTGTTATTGGCCCATTACTGTTTGCTGTTATTTATAATCATTCACTACCAATTTG GGATGGCTGGATTTGGATTATTGGTTTAGCGTTTTACTGTATTATTATCCTGCTATCGATGA CCTTCATGTTAACCCCTCAAGCTCAGGGGAGTAAACAGGAGACAAGTGCTTAGTTATTTCG TCACCAAATGATGTTATTCCGCGAAATATAATGACCCTCTTGATAACCCAAGAGCATCACA |

TATACCTGCCGTTCACTATTATTTAGTGAAATGAGATATTATGATATTTTCTGAATTGTGAT
TAAAAAGGCAACTTTATGCCCATGCAACAGAAACTATAAAAAATACAGAGAATGAAAAG
AAACAGATAGATTTTTTAGTTCTTTAGGCCCGTAGTCTGCAAATCCTTTTATGATTTTCTAT
CAAACAAAAGAGGAAAATAGACCAGTTGCAATCCAAACGAGAGTCTAATAGAATGAGGT
CGAAAAGTAAATCGCGCGGGTTTGTTACTGATAAAGCAGGCAAGACCTAAAATGTGTAAA
GGGCAAAGTGTATACTTTGGCGTCACCCCTTACATATTTTAGGTCTTTTTTTATTGTGCGTA
ACTAACTTGCCATCTTCAAACAGGAGGGCTGGAAGAAGCAGACCGCTAACACAGTACATA
AAAAAGGAGACATGAACGATGAACATCAAAAAGTTTGCAAAACAAGCAACAGTATTAAC
CTTTACTACCGCACTGCTGGCAGGAGGCGCAACTCAAGCGTTTGCGAAAGAAACGAACCA
AAAGCCATATAAGGAAACATACGGCATTTCCCATATTACACGCCATGATATGCTGCAAAT
CCCTGAACAGCAAAAAAATGAAAAATATCAAGTTCCTGAGTTCGATTCGTCCACAATTAA
AAATATCTCTTCTGCAAAAGGCCTGGACGTTTGGGACAGCTGGCCATTACAAAACGCTGA
CGGCACTGTCGCAAACTATCACGGCTACCACATCGTCTTTGCATTAGCCGGAGATCCTAAA
AATGCGGATGACACATCGATTTACATGTTCTATCAAAAAGTCGGCGAAACTTCTATTGACA
GCTGGAAAAACGCTGGCCGCGTCTTTAAAGACAGCGACAAATTCGATGCAAATGATTCTA
TCCTAAAAGACCAAACACAAGAATGGTCAGGTTCAGCCACATTTACATCTGACGGAAAAA
TCCGTTTATTCTACACTGATTTCTCCGGTAAACATTACGGCAAACAAACACTGACAACTGC
ACAAGTTAACGTATCAGCATCAGACAGCTCTTTGAACATCAACGGTGTAGAGGATTATAA
ATCAATCTTTGACGGTGACGGAAAAACGTATCAAAATGTACAGCAGTTCATCGATGAAGG
CAACTACAGCTCAGGCGACAACCATACGCTGAGAGATCCTCACTACGTAGAAGATAAAGG
CCACAAATACTTAGTATTTGAAGCAAACACTGGAACTGAAGATGGCTACCAAGGCGAAGA
ATCTTTATTTAACAAAGCATACTATGGCAAAAGCACATCATTCTTCCGTCAAGAAAGTCAA
AAACTTCTGCAAAGCGATAAAAAACGCACGGCTGAGTTAGCAAACGGCGCTCTCGGTATG
ATTGAGCTAAACGATGATTACACACTGAAAAAAGTGATGAAACCGCTGATTGCATCTAAC
ACAGTAACAGATGAAATTGAACGCGCGAACGTCTTTAAAATGAACGGCAAATGGTACCTG
TTCACTGACTCCCGCGGATCAAAAATGACGATTGACGGCATTACGTCTAACGATATTTACA
TGCTTGGTTATGTTTCTAATTCTTTAACTGGCCCATACAAGCCGCTGAACAAAACTGGCCTT
GTGTTAAAAATGGATCTTGATCCTAACGATGTAACCTTTACTTACTCACACTTCGCTGTACC
TCAAGCGAAAGGAAACAATGTCGTGATTACAAGCTATATGACAAACAGAGGATTCTACGC
AGACAAACAATCAACGTTTGCGCCAAGCTTCCTGCTGAACATCAAAGGCAAGAAAACATC
TGTTGTCAAAGACAGCATCCTTGAACAAGGACAATTAACAGTTAACAAATAAAAACGCAA
AAGAAAATGCCGATATTGACTACCGGAAGCAGTGTGACCGTGTGCTTCTCAAATGCCTGA
TTCAGGCTGTCTATGTGTGACTGTTGAGCTGTAACAAGTTGTCTCAGGTGTTCAATTTCATG
TTCTAGTTGCTTTGTTTTACTGGTTTCACCTGTTCTATTAGGTGTTACATGCTGTTCATCTGT
TACATTGTCGATCTGTTCATGGTGAACAGCTTTAAATGCACCAAAAACTCGTAAAAGCTCT
GATGTATCTATCTTTTTTACACCGTTTTCATCTGTGCATATGGACAGTTTTCCCTTTGAT

**recA1::ampR**

CCATCTCTACCGGTTCGCTTTCACTGGATATCGCGCTTGGGGCAGGTGGTCTGCCGATGGGCCGTAT
CGTCGAAATCTACGGACCGGAATCTTCCGGTAAAACCACGCTGACGCTGCAGGTGATCGCCGCAGC
GCAGCGTGAAGGTAAAACCTGTGCGTTTATCGATGCTGAACACGCGCTGGACCCAATCTACGCACG
TAAACTGGGCGTCGATATCGACAACCTGCTGTGCTCCCAGCCGGACACCGGCGAGCAGGCACTGGA

AATCTGTGACGCCCTGGCGCGTTCTGGCGCAGTAGACGTTATCGTCGTTGACTCCGTGGCGGCACTG
ACGCCGAAAGCGGAAATCGAAGGCGAAATC***GAT***GACTCTCACATGGGCCTTGCGGCACGTATGATG
AGCCAGGCGATGCGTAAGCTGGCGGGTAACCTGAAGCAGTCCAACACGCTGCTGATCTTCATCAAC
CAGATCCGTATGAAAATTGGTGTGATGTTCGGTAACCCGGAAACCACTACCGGTGGTAACGCGCTG
AAATTCTACGCCTCTGTTCGTCTCGACATCCGTCGTATCGGCGCGGTGAAAGAGGGCGAAAACGTG
GTGGGTAGCGAAACCCGCGTGAAAGTGGTGAAGAACAAAATCGCTGCGCCGTTTAAACAGGCTGAA
TTCCAGATCCTCTACGGCGAAGGTATCAACTTCTACGGCGAACTGGTTGACCTGGGCGTAAAAGAG
AAGCTGATCGAGAAAGCAGGCGCGTGGTACAGCTACAAAGGTGAGAAGATCGGTCAGGGTAAAGC
GAATGCGACTGCCTGGCTGAAAGATAACCCGGAAACCGCGAAAGAGATCGAGAAGAAAGTACGTG
AGTTGCTGCTGAGCAACCCGAACTCAACGCCGGATTTCTCTGTAGATGATAGCGAAGGCGTAGCAG
AAACTAACGAAGATTTTTAATGATTGCAGTCCAGTTACGCTGGAGTCTGAGGCTCGTCCTGAATGAT
ATCAAGCTTGAATTCGTTGGTTCATCCCGTGGGCATTGCATAGGGATAACAGGGTAATCTAAATACA
TTCAAATATCTATCCGCTCATGAGACAATAACCCTGATAAATGCTTCAATAATATTGAAAAAGGAAG
AATATGAGTATTCAACATTTCCGTGTCGCCCTTATTCCCTTTTTTGCGGCATTTTGCCTTCCTGTTTTT
GCTCACCCAGAAACGCTGGTGAAAGTAAAAGATGCCGAAGATCAGTTGGGTGCACGTGTGGGTTAC
ATCGAACTGGACCTCAACAGCGGTAAGATTCTTGAGAGTTTTCGCCCCGAAGAACGTTTCCCAATGA
TGAGCACTTTTAAAGTTCTGCTCTGTGGCGCGGTATTATCCCGTATTGACGCCGGGCAAGAGCAACT
CGGTCGCCGCATACACTATTCTCAGAATGACTTGGTTGAGTACTCACCAGTCACAGAAAAGCATCTT
ACGGACGGCATGACAGTACGCGAATTATGCAGCGCTGCCATAACCATGAGTGATAACACGGCGGCC
AACTTACTTCTGACAACGATCGGAGGACCGAAGGAGCTTACCGCTTTTTTGCACAACATGGGTGATC
ATGTAACTCGCCTTGATCGTTGGGAACCGGAGCTGAATGAAGCCATACCAAACGACGAGCGTGACA
CCACGATGCCTGTAGCTATGGCAACAACGTTGCGCAAACTCTTAACTGGCGAACTTCTTACTCTCGC
TTCCCGGCAACAATTAATAGACTGGATGGAGGCGGATAAAGTTGCAGGACCACTTCTGCGCTCGGC
CCTTCCGGCTGGCTGGTTTATTGCTGATAAATCTGGAGCCGGTGAGCGTGGGTCCCGCGGTATTATT
GCAGCCCTGGGGCCAGATGGTAAGCCCTCCCGTATCGTAGTTATCTACACGACGGGGAGCCAGGCA
ACTATGGACGAACGTAATCGCCAGATCGCTGAGATAGGTGCCTCCCTGATTAAGCATTGGTAATAA
CCAGGCATCTCGTCTTGTTTGATACACAAGGGTCGCATCTGCGGCCCTTTTGCTTTTTTAAGTTGTAA
GGATATGCCATGACAGAATCAACATCCCGTCGCCCGGCATATGCTCGCCTGTTGGATCGTGCGGTAC
GCATTCTGGCGGTGCGCGATCACAGTGAGCAAGAACTGCGACGTAAACTCGCGGCACCGATTATGG
GCAAAAATGGCCCAGAAGAGATTGATGCTACGGCAGAAGATTACGAGCGCGTTATTG

**ΔsbcD::ampR**

TGATTGCAGTCCAGTTACGCTGGAGTCTGAGGCTCGTCCTGAATGATATCAAGCTTGAATTCGTTGG
TTCATCCCGTGGGCATTGCATAGGGATAACAGGGTAATCTAAATACATTCAAATATCTATCCGCTCA
TGAGACAATAACCCTGATAAATGCTTCAATAATATTGAAAAAGGAAGAATATGAGTATTCAACATT
TCCGTGTCGCCCTTATTCCCTTTTTTGCGGCATTTTGCCTTCCTGTTTTTGCTCACCCAGAAACGCTGG
TGAAAGTAAAAGATGCCGAAGATCAGTTGGGTGCACGTGTGGGTTACATCGAACTGGACCTCAACA
GCGGTAAGATTCTTGAGAGTTTTCGCCCCGAAGAACGTTTCCCAATGATGAGCACTTTTAAAGTTCT
GCTCTGTGGCGCGGTATTATCCCGTATTGACGCCGGGCAAGAGCAACTCGGTCGCCGCATACACTAT
TCTCAGAATGACTTGGTTGAGTACTCACCAGTCACAGAAAAGCATCTTACGGACGGCATGACAGTA
CGCGAATTATGCAGCGCTGCCATAACCATGAGTGATAACACGGCGGCCAACTTACTTCTGACAACG
ATCGGAGGACCGAAGGAGCTTACCGCTTTTTTGCACAACATGGGTGATCATGTAACTCGCCTTGATC
GTTGGGAACCGGAGCTGAATGAAGCCATACCAAACGACGAGCGTGACACCACGATGCCTGTAGCTA
TGGCAACAACGTTGCGCAAACTCTTAACTGGCGAACTTCTTACTCTCGCTTCCCGGCAACAATTAAT
AGACTGGATGGAGGCGGATAAAGTTGCAGGACCACTTCTGCGCTCGGCCCTTCCGGCTGGCTGGTTT
ATTGCTGATAAATCTGGAGCCGGTGAGCGTGGGTCCCGCGGTATTATTGCAGCCCTGGGGCCAGAT

GGTAAGCCCTCCCGTATCGTAGTTATCTACACGACGGGGAGCCAGGCAACTATGGACGAACGTAAT
CGCCAGATCGCTGAGATAGGTGCCTCCCTGATTAAGCATTGGTAATAACCAGGCAT

**del_sbcD_p1**

gatttccgtggcgcagaaaaaagcaaatggcacatctgtttgggtataatcgcgcccatgctttttcgccaTGATTGCAGTCCAGTTACG

**del_sbcD_p2**

gggtgaaatcaatcttccattcgccttttaatgagttcaggtttttcaggcgcaggctgagaattATGCCTGGTTATTACCAATGCTTAA

**Δcas1::purR**

GATCATTTCACGAATTTTTGCGGATACATCACCTACATATACCCCTGCACGTACCTCCAACAACCAG
ATGGCTAATCTGCCTCGTAAGCGCGGAGGTACATTTTCAGTGACCACGACCAACATACTCATTTCCT
GACGGATGGCCTTTTTGCGTTTCTACAAACTCTTTTTGTTTATTTTTCTAAATACATTCAAATATGTAT
CCGCTCATGAGACAATAACCCTGATAAATGCTTCAATAATATTGAAAAAGGAAGAGTATGACTGAA
TACAAGCCCACGGTACGCTTGGCGACGCGCGACGATGTTCCCCGCGCTGTTCGTACATTAGCTGCGG
CCTTTGCAGATTACCCAGCGACGCGCCATACGGTCGATCCGGACCGCCATATCGAGCGTGTCACAG
AATTGCAGGAACTTTTCTTAACTCGCGTGGGCCTTGACATCGGAAAGGTCTGGGTGGCTGACGATGG
CGCTGCAGTGGCTGTTTGGACCACTCCGGAGAGTGTAGAGGCTGGTGCAGTGTTCGCCGAAATTGGT
CCTCGTATGGCCGAATTAAGTGGAAGTCGTCTGGCAGCCCAACAACAAATGGAAGGGTTGCTTGCG
CCCCACCGTCCGAAAGAACCCGCGTGGTTCCTTGCCACCGTTGGAGTAAGCCCAGATCACCAGGGG
AAGGGTTTAGGATCTGCCGTAGTTTTACCAGGTGTGGAGGCAGCAGAACGTGCGGGAGTTCCGGCC
TTCCTTGAGACGTCGGCGCCGCGCAATTTACCGTTTTACGAACGTCTTGGATTCACCGTTACGGCGG
ACGTGGAGGTGCCGGAGGGACCCCGTACTTGGTGTATGACTCGTAAACCGGGAGCCTGATAATTTA
TTACACCTCAATCACAGTGGAGCCAAAGATAGCAAGCCACATCCCATCGATTTAGCTGGCCCAATA
CCTTGCTGTACAAGATCTATTAACGCTGGCGCGTCGTTGATGGTGAGCACACCTTCAAAGCAAACCG

**Δcas2::purR**

GCGGTTTATCCCCGCTGGCGCGGGGAACTCTCTAAAAGTATACATTTGTTCTTAAAGCATTTTTTCCC
ATAAAAACAACCCACCAACCTTAATGTAACATTTCCTTATTATTAAAGATCAGCTAATTCTTTGTTTT
CCTGACGGATGGCCTTTTTGCGTTTCTACAAACTCTTTTTGTTTATTTTTCTAAATACATTCAAATATG
TATCCGCTCATGAGACAATAACCCTGATAAATGCTTCAATAATATTGAAAAAGGAAGAGTATGACT
GAATACAAGCCCACGGTACGCTTGGCGACGCGCGACGATGTTCCCCGCGCTGTTCGTACATTAGCTG
CGGCCTTTGCAGATTACCCAGCGACGCGCCATACGGTCGATCCGGACCGCCATATCGAGCGTGTCAC
AGAATTGCAGGAACTTTTCTTAACTCGCGTGGGCCTTGACATCGGAAAGGTCTGGGTGGCTGACGAT
GGCGCTGCAGTGGCTGTTTGGACCACTCCGGAGAGTGTAGAGGCTGGTGCAGTGTTCGCCGAAATT
GGTCCTCGTATGGCCGAATTAAGTGGAAGTCGTCTGGCAGCCCAACAACAAATGGAAGGGTTGCTT
GCGCCCCACCGTCCGAAAGAACCCGCGTGGTTCCTTGCCACCGTTGGAGTAAGCCCAGATCACCAG
GGGAAGGGTTTAGGATCTGCCGTAGTTTTACCAGGTGTGGAGGCAGCAGAACGTGCGGGAGTTCCG
GCCTTCCTTGAGACGTCGGCGCCGCGCAATTTACCGTTTTACGAACGTCTTGGATTCACCGTTACGG
CGGACGTGGAGGTGCCGGAGGGACCCCGTACTTGGTGTATGACTCGTAAACCGGGAGCCTGATAAT
TCAGCTACTCCGATGGCCTGCATCTCCCAGTGAAACAGGAAGCGGAATGGCAACAGGCTGTGCATC
TTCAGGTGGGGCCGGCGGTTGTATTTCTCCAGCGGCAAGCACGTCCTCTATAAGCGGAATCAATT

| |
|---|
| **Δcas3::Term-ugpBp-sspB-TZ yibDp-casA** |
| TATGAGCAGCATCGAAAAATAGCCCGCTGATATCATCGATAATACTAAAAAAACAGGGAGGCTATT<br>ACCAGGCATCAAATAAAACGAAAGGCTCAGTCGAAAGACTGGGCCTTTCGTTTTATCTGTTGTTTGT<br>CGGTGAACGCTCTCTACTAGAGTCACACTGGCTCACCTTCGGGTGGGCCTTTCTGCGTTTATATCTTT<br>CTGACACCTTACTATCTTACAAATGTAACAAAAAAGTTATTTTTCTGTAATTCGAGCATGTCATGTTA<br>CCCCGCGAGCATAAAACGCGTGTGTAGGAGGATAATCTATGGATTTGTCACAGCTAACACCACGTC<br>GTCCCTATCTGCTGCGTGCATTCTATGAGTGGTTGCTGGATAACCAGCTCACGCCGCACCTGGTGGT<br>GGATGTGACGCTCCCTGGCGTGCAGGTTCCTATGGAATATGCGCGTGACGGGCAAATCGTACTCAA<br>CATTGCGCCGCGTGCTGTCGGCAATCTGGAACTGGCGAATGATGAGGTGCGCTTTAACGCGCGCTTT<br>GGTGGCATTCCGCGTCAGGTTTCTGTGCCGCTGGCTGCCGTGCTGGCTATCTACGCCCGTGAAAATG<br>GCGCAGGCACGATGTTTGAGCCTGAAGCTGCCTACGATGAAGATACCAGCATCATGAATGATGAAG<br>AGGCATCGGCAGACAACGAAACCGTTATGTCGGTTATTGATGGCGACAAGCCAGATCACGATGATG<br>ACACTCATCCTGACGATGAACCTCCGCAGCCACCACGCGGTGGTCGACCGGCATTACGCGTTGTGA<br>AGTAATCGACCTAGCATAACCCCGCGGGGCCTCTTCGGGGGTCTCGCGGGGTTTTTTGCTGAAAGAA<br>GCTTCAAATAAAACGAAAGGCTCAGTCGAAAGACTGGGCCTTTCGTTTTATCTGTTGTTTGTCGCTG<br>CGGCCGGGTCAGGTATGATTTAAATGGTCAGTAACGGGTCTTGAGGGGTTTTTTGCCACAGCTAACA<br>CCACGTCGTCCCTATCTGCTGCCCTAGGTCTATGAGTGGTTGCTGGATAACGTGCGTAATTGTGCTG<br>ATCTCTTATATAGCTGCTCTCATTATCTCTCTACCCTGAAGTGACTCTCTCACCTGTAAAAATAATAT<br>CTCACAGGCTTAATAGTTTCTTAATACAAAGCCTGTAAAACGTCAGGATAACTTCTATATTCAGGGA<br>GACCACAACGGTTTCCCTCTACAAATAATTTTGTTTAACTTTTGAAGGAGAACAAATGAATTTGCTT<br>ATTGATAACTGGATCCCTGTACGCCCGCGAAACGGGGGGAAAGTCCAAATCATAAATCTGCAATCG<br>CTATACTGCAGTAGAGATCAGTGGCGATTAAGTTTGCCCCGTGACGATATGGAACTGGCCGCTTTAG<br>CACTGCTGGTTTGCATTGGGCAAATTATCGCCCCGGCAAAAGATGACGTTGAATTTCGACATCGCAT<br>AATGAATCCGCTCACTGAAGATGAGT |
